# Modularity in temporal co-occurrence networks at a stopover site reveals strong co-migration fidelity in African-European migratory landbirds over time

**DOI:** 10.1101/2022.02.17.480723

**Authors:** Bruno Bellisario, Massimiliano Cardinale, Ivan Maggini, Leonida Fusani, Claudio Carere

## Abstract

Migratory species are changing the timing of departures from wintering areas and arrival to breeding sites (i.e., migration phenology), as response to climate change to exploit maximum food availability at higher latitudes and improve their fitness. Despite the impact of changing migration phenology on migratory connectivity at population and community level, the extent to which such individual and species-specific response affects associations among co-migrating species has been seldom explored. By applying temporal co-occurrence network models on 15 years of standardized bird ringing data at a spring stopover site, we show that African-European migratory landbirds tend to migrate in well-defined groups of species with high temporal overlap. Such ‘co-migration fidelity’ is significantly increasing over the years and higher in trans-Saharan than North-African migrants. Our findings suggest non-random patterns of associations in co-migrating species, possibly related to the existence of regulatory mechanisms associated with changing climate conditions and different uses of stopover sites by migrants, ultimately influencing the global migration economy of landbirds in the Palearctic-African migration system.

## 1. Introduction

Migration represents a key life-history event for a large number of species and is typically associated with the temporal and geographic variation in resource availability [1]. Alongside endogenous (e.g., circannual rhythms) and exogenous (e.g., photoperiodic cues) mechanisms [2,3], favourable environmental conditions and food availability *en route* are key elements driving the timing of migration in birds [4–6].

Climate change may pose serious problems to migrants spending different parts of their annual cycle in different parts of the world as it drives the restructuring of important phenological events [7–9]. For instance, arrival dates to breeding grounds of Central-European populations of Pied flycatchers (*Ficedula hypoleuca*) showed no significant synchronization with advances in food-peak dates due to warming temperature, leading to a drastic decline (ca. 90%) in the abundance of populations that arrived after the peak [10]. Overall, differences in the extent of climatic change along migratory routes, as well as at wintering, stopover and breeding areas [11–13], increase the chance of phenological mismatches, influencing the interactions among co-occurring species including their food sources, predators, and competitors.

Within this context, migratory connectivity, defined as the ‘geographic linking of individuals and populations between one life cycle stage and another’ [14] represents a key factor having far-reaching consequences at multiple ecological scales, from individual fitness to community dynamics and ecosystem functioning. To date, most studies rely on a purely spatial perspective aimed at determining the extent to which different areas of a species’ annual range are linked by the movement paths of individuals [14,15], providing useful insights on how climate change influence the ability of such areas in sustaining migratory flows (e.g., ‘migration hot spots’ [16]). However, an explicit consideration of the temporal dimension [17] is fundamental to understand how, when and for how long species change their migratory patterns and what are the consequences at multiple ecological levels.

Advances in the date of departure from wintering areas in response to endogenous and/or exogenous factors reflecting changing climate conditions have been observed in several species [18–20]. Populations of origin, sex, age, ecological requirements might all affect the ability of species to change migration phenology [21–26], leading in some cases to asynchronous migration patterns. Moreover, phenotypic plasticity and microevolutionary changes can influence the long-term responses of species to track changes along migratory routes, leading in this case to differences in the consistency of migration pattern (i.e., how repeatable migration phenology and synchrony are over time [17]).

Migrations involve the simultaneous movement of multiple species at once, connecting separated and diverse communities that may have a strong impact on the dynamic and stability of ecosystems. Migratory routes of different species often converge both in space and time [27], resulting in direct and indirect ecological interactions such as competition for resources, predation and even social interactions. For instance, studies have shown how parasites transmission, as well as transient dynamics in lake ecosystems, can be related to migration timing [28,29]. Similarly, intra- and interspecific competition may be affected by synchronous migrations by excluding inferior competitors from high-quality foraging areas [30,31], or promoting coexistence when the joint use of a resource increases its quality and productivity [32]. Migration timing can rely on social interactions, as observed in songbirds migrating along the coasts of Baltic Sea, which use vocalizations from conspecifics as cues to assess the suitability of potential stopover habitats [33,34]. However, despite the ecological importance of interactions between co-migrants, we still lack a proper understanding on how long-term phenological changes affect the restructuring of avian assemblages *en route*.

Investigating the causes and the effects of changing migration phenology from a community point of view entails inherent difficulties associated with the need to constantly track and monitor animals across oceans and continents [27]. A further source of uncertainty derives from the need to appropriately model multi-species temporal movements to identify recurrent patterns in the aggregation of species and forecast the consequences of climate change on migrations. Within this context, co-occurrence analysis and co-occurrence networks are valuable tools to elucidate mechanisms driving species assemblages. Recently, García-Navas et al. (2021) [35] used null model approaches on 20 years of abundance data to investigate co-occurrence patterns in bird communities inhabiting habitats with different degree of productivity, showing the role of social information and microhabitat preferences over interspecific competition. Furthermore, Montaño-Centellas (2020) [36] used spatial data to build and explore co-occurrence networks of avian mixed-species flocks across elevation gradients, identifying changes in interactions as the main driver of species turnover among Andean flocking birds. Co-occurrence networks can thus help illustrating complex properties of animal movements with simple graphical metrics [37], although the actual ecological meaning of revealed links must be critically evaluated (see [38] and references therein).

By using temporal co-occurrence network models on time series data spanning 15 years of standardized bird ringing activity at the Mediterranean spring stopover area of Ponza (Italy), we searched for a consistent and repeated order of association over time to study how long-term changes in the migration timing of African-European migratory landbirds affect the structuring of associations among co-migrants.

## 2. Materials and methods

### 2.1 Sampling site and bird ringing activity

The study was conducted on Ponza (figure 1), a small island in the Tyrrhenian Sea with an extension of ca. 9.87 km^2^, located about 50 km off the Western coast of Italy (40°55’ N, 12°58’ E), where we have been monitoring spring migration through capture and ringing since 2002 (http://www.inanellamentoponza.it). Ponza is located along one of the main Mediterranean migratory routes and attracts a large number of African-European migratory landbirds during spring migration [20]. The bird capture season started in March and ended in May almost every year (exact start and end dates can be found in the supplementary Table S1 in [20]). Mist nets were opened every day except for days with heavy rain or strong winds (>15 knots) and the number of days with reduced effort was less than 1% of the total. Mist-nets were checked hourly from dawn until one hour after dusk, and birds were captured throughout the day with an average of 227 m of mist-nets over the time-period. We kept the net brand and model (Lavorazione Reti Bonardi, Monte Isola, BS, Italy, http://www.vbonardi.it/; 2.4 m height, 16 mm mesh size), as well as the vegetation height constant throughout the entire study period. After being captured, birds were ringed and measured using standard EURING procedures [39]. Here we used the daily abundances of the 31 most abundant species passing through Ponza (table 1), using ringing data over a period of 15 years (2007 to 2021), during which the procedures were fully standardized.

**Figure 1.**
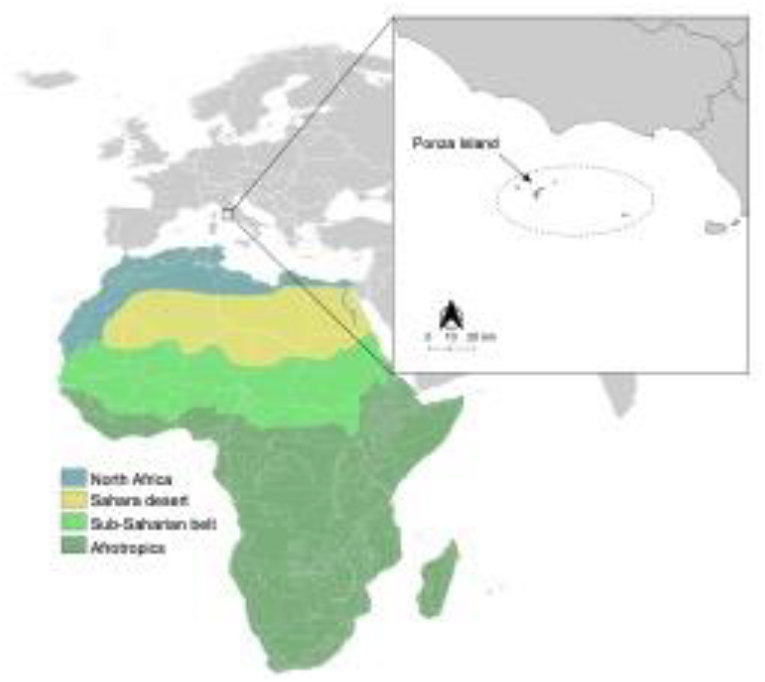
Geographical framework of the study area within the Palearctic-African migration system. The dotted circle in the box shows the ‘Pontine’ islands system with the exact location of Ponza Island. Colours in the map show a synthetic view of the main African ecoregions corresponding to the supposed wintering areas of species analysed in this study.

**Table 1.** List of the 31 most abundant species ringed at the stop over site of Ponza Island during the period 2007-2021.

| Species | Wintering area | Foraging niche | Number of sampled individuals |  |  |
| --- | --- | --- | --- | --- | --- |
| | | | Total | Mean (± SD) | Range [min-max] |
| *Acrocephalus arundinaceus* | Tropical | Invertivore glean arboreal | 1,156 | 25.2 ± 11.391 | [13-58] |
| *Acrocephalus schoenobaenus* | Sahel | Invertivore glean arboreal | 378 | 77.067±48.792 | [40-236] |
| *Anthus trivialis* | Tropical | Invertivore ground | 2,197 | 146.467±44.171 | [72-207] |
| *Delichon urbica* | Tropical | Invertivore sally air | 1,283 | 85.534±49.393 | [21-187] |
| *Erithacus rubecula* | North Africa | Invertivore ground | 14,948 | 996.534±877.128 | [84-2599] |
| *Ficedula albicollis* | Tropical | Invertivore sally air | 11,504 | 121.734±99.349 | [5-279] |
| *Ficedula hypoleuca* | Tropical | Invertivore sally air | 1,826 | 766.934±271.304 | [369-1393] |
| *Fringilla coelebs* | Tropical | Invertivore ground | 421 | 28.067±20.537 | [1-58] |
| *Hippolais icterina* | Tropical | Invertivore glean arboreal | 18,850 | 1256.667±706.695 | [325-2922] |
| *Hirundo rustica* | Sahel | Invertivore sally air | 6,402 | 426.8±132.102 | [208-700] |
| *Jynx torquilla* | Sahel | Invertivore ground | 501 | 33.4±15.263 | [17-67] |
| *Lanius senator* | Tropical | Invertivore ground | 621 | 41.4±14.603 | [17-71] |
| *Luscinia megarhynchos* | Tropical | Invertivore ground | 1,976 | 131.733±50.941 | [56-214] |
| *Merops apiaster* | Sahel | Invertivore sally air | 757 | 50.466±26.848 | [21-114] |
| *Muscicapa striata* | Sahel | Invertivore sally air | 9,500 | 633.333±246.003 | [187-1108] |
| *Oenanthe oenanthe* | Sahel | Invertivore sally ground | 2,222 | 148.133±59.677 | [67-259] |
| *Oriolus oriolus* | Tropical | Granivore ground | 1,343 | 89.533±37.557 | [39-182] |
| *Phoenicurus ochruros* | North Africa | Invertivore sally ground | 2,750 | 183.333±142.749 | [6-397] |
| *Phoenicurus phoenicurus* | Sahel | Invertivore sally ground | 6,728 | 448.533±255.998 | [155-1118] |
| *Phylloscopus collybita* | North Africa | Invertivore glean arboreal | 5,880 | 392±288.111 | [88-1050] |
| *Phylloscopus sibilatrix* | Tropical | Invertivore glean arboreal | 15,402 | 1026.8±285.567 | [597-1430] |
| *Phylloscopus trochilus* | Sahel | Invertivore glean arboreal | 13,466 | 897.733±315.275 | [424-1421] |
| *Saxicola rubetra* | Tropical | Invertivore sally ground | 13,392 | 892.8±432.842 | [7-1646] |
| *Saxicola torquata* | North Africa | Invertivore sally ground | 780 | 52±60.523 | [2-184] |
| *Streptopelia turtur* | Sahel | Granivore ground | 724 | 48.266±36.526 | [18-139] |
| *Sylvia atricapilla* | North Africa | Invertivore glean arboreal | 13,885 | 925.666±1738.913 | [29-5614] |
| *Sylvia borin* | Tropical | Invertivore glean arboreal | 35,655 | 2377±1485.051 | [337-5967] |
| *Sylvia cantillans* | Sahel | Invertivore glean arboreal | 14,117 | 941.133±659.287 | [259-2789] |
| *Sylvia communis* | Sahel | Invertivore glean arboreal | 25,358 | 1690.533±1023.444 | [40-3589] |
| *Turdus philomelos* | North Africa | Invertivore ground | 1,985 | 132.333±146.932 | [16-477] |
| *Upupa epops* | Sahel | Invertivore ground | 526 | 35.066±18.525 | [15-76] |

### 2.2 Network inference

For each year of survey (*n* = 15), we reconstructed the temporal co-occurrence network by calculating the time-lagged Spearman rank correlations using the daily abundances of each species with each other in the previous time step (i.e., the lag-1 correlation [40]). To avoid spurious correlations due to extremely low abundances, we removed from computation those species for which the total number of individuals in each year was below the 25^th^ percentile of the entire distribution. We used the packages ‘Hmisc’ [41] and ‘qvalue’ [42] of R [43] to adjust the significance of correlations using the Bonferroni-Holm procedure. Co-occurrences were considered significant for correlation coefficients *r* > 0.5 and *p* < 0.05. We used unweighted links, where potential co-occurrences were considered irrespective of the strength of the correlation.

### 2.3 Network modularity

We used an eigenvector-based maximizing algorithm (i.e., the Louvain algorithm [44]) to measure modularity (*Q*), that is, the degree to which a network subdivides in densely connected groups of nodes (aka modules) with only sparser connections between groups [45]. From an ecological point of view, modularity is a straightforward structural property able to detect groups of species having similar responses to the main processes structuring biodiversity in space and time [46–48]. In our case, the higher the modularity, the greater will be the tendency of networks to cluster into subgroups of species characterized by common migration timing (i.e., high temporal co-occurrence). We considered networks with *Q* > 0.3 as having a strong modular subdivision [45].

The yearly measured modularity values were filtered by means of a simple moving average (SMA), to reduce background noise in the original time series and emphasize trends. We used SMA with an automated selection procedure based on the Akaike information criterion to find the optimal order of the moving average [49]. Modularity was measured by using the function ‘cluster_louvain’ [44] in the package ‘igraph’ [50] of R, while we used the functions ‘sma’ and ‘sen.slope’ in the packages ‘smooth’ [49] and ‘trend’ [51], respectively, to smooth the time series and measure the Mann-Kendall statistics.

### 2.4 Co-migration fidelity

We defined ‘co-migration fidelity’ as the frequency with which two species tended to co-occur in the same module over the years, measured by using the normalized mutual information (*nmi* [52]). Basically, this metric assesses whether two classifications (i.e., module affiliation in different years) on the same set (i.e., species) well explain one another, providing a robust metric for comparing networks as it is not affected by the number and size of modules found in each network. The normalized mutual information is bounded between 0 and 1, so that if two species were always part of the same module over the years (i.e., perfect congruence, high fidelity), then *nmi* = 1; conversely, if two species never co-occurred within the same module over the years (i.e., independent, no fidelity), then *nmi* = 0. We thus measured all *nmi* values between species, resulting in a symmetric similarity matrix expressing the pairwise mutual information among species.

We finally used the permutational multivariate analysis of variance using distance matrices for partitioning the measured normalized mutual information among sources of variation given by wintering areas and foraging niches. We therefore divided the species in three main groups (table 1) corresponding to those wintering either north of the Sahara Desert (North African), in the sub-Saharan belt extended between the Sahara Desert and the Sudan savannah (Sahel), or in the Afrotropical ecozone (Tropical). Following Pigot et al. (2020) [53], species were also characterized by considering their main foraging niches (table 1). The normalized mutual information and permutational multivariate analysis of variance were measured by using the function ‘compare’ and ‘adonis2’ in the ‘igraph’ and ‘vegan’ [50,54] packages of R, respectively.

## 3. Results

A total of 226,533 individuals were ringed over the 15 years of field activity, with an average (± SD) of 15,117 ± 4,031 per year and a minimum of 8,861 in 2010 and a maximum of 23,186 in 2015. Three main families accounted for 84% of the total, Sylviidae (four species, 40%), Muscicapidae (seven species, 29%) and Phylloscopidae (three species, 15 %). The garden warbler (*Sylvia borin*) and common whitethroat (*Sylvia communis*) were the most abundant species, while other as the common chaffinch (*Fringilla coelebs*) and the great reed warbler (*Acrocephalus arundinaceus*) were less abundant (table 1).

Overall, temporal co-occurrence networks always showed relatively strong modular structures, with an average *Q* (± SD) of 0.357 ± 0.053 and ranging from a minimum of 0.288 in 2010 and a maximum of 0.448 in 2018, also showing a significant increase over the years (Mann-Kendall trend test *z* = 3.865,*p* < 0.001) (figure 2). The pairwise co-migration fidelity, measured by the normalized mutual information (*nmi*), showed a well-defined pattern, with species subdividing in (at least) six to seven main clusters (figure 3a). Interestingly, one main cluster was characterized by species with very high levels of co-migration fidelity (*S. borin*, *Hippolais icterina*, *Muscicapa striata*, *Oriolus oriolus*), and always co-occurring in the same module with the same species over the years, with no temporal overlap with other species (figure 3a). Other, as the black redstart (*Phoenicurus ochruros*), the song thrush (*Turdus philomelos*), the common stonechat (*Saxicola torquata*), the European robin (*Erithacus rubecula*) and the common chiffchaff (*Phylloscopus collybita*) showed more variable *nmi* index, with partial temporal overlap with other species (figure 3a). Co-migration fidelity significantly differed between wintering areas (*F* = 2.635, *p* = 0.004), with birds wintering in North Africa showing on average lower values of *nmi* compared to those wintering in the sub-Saharan belt and Afrotropical ecozone (figure 3b). We did not find significant differences with respect to foraging niches (*F* = 1.379, *p* = 0.088, figure 3b).

**Figure 2.**
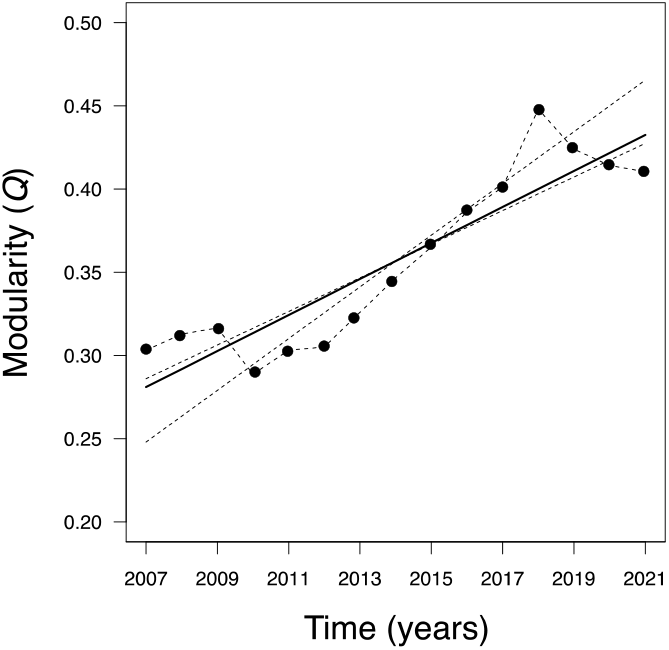
Trend of modularity (*Q*) in temporal co-occurrence networks. Dashed black lines indicate the original time series, while solid and dotted lines show the measured trends and uncertainty in trend slopes, respectively.

**Figure 3.**
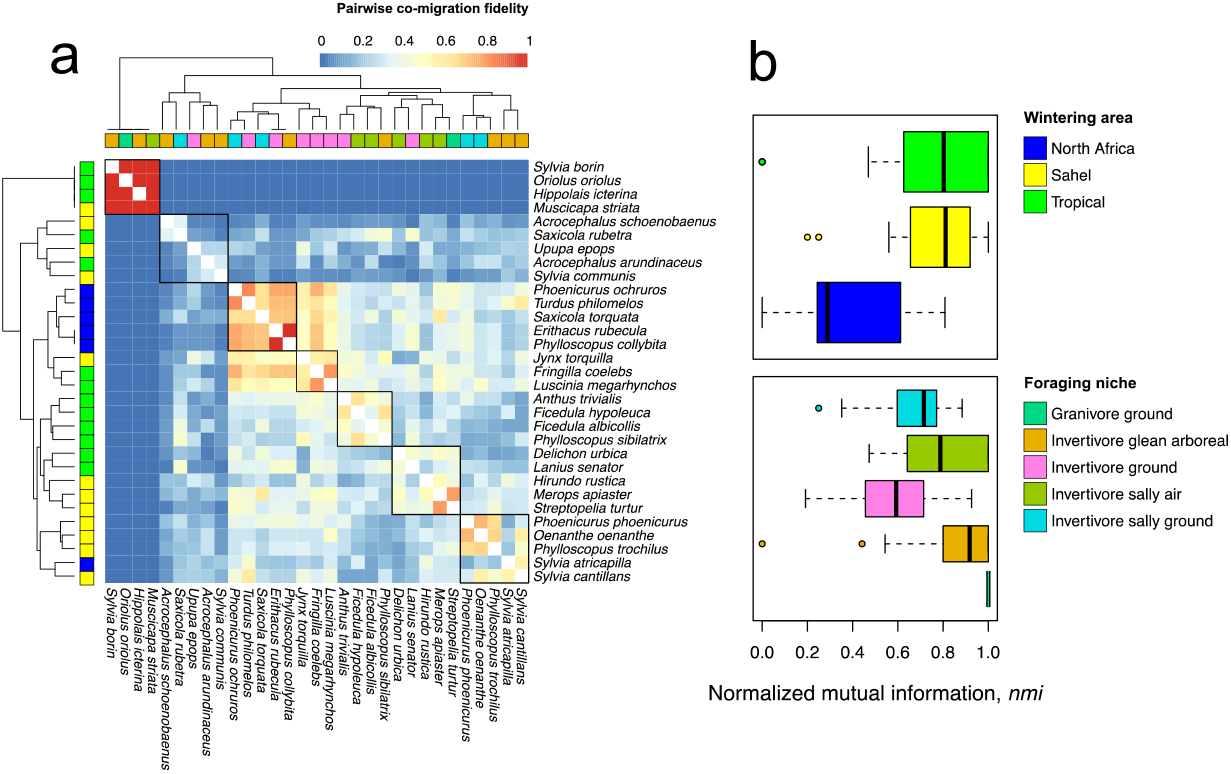
(a) Heatmap showing the species’ pairwise co-migration fidelity measured by the normalized mutual information, *nmi*. Black boxes within the heatmap correspond to the main cluster subdivision, while coloured boxes below and alongside dendrograms show to which foraging niche and wintering area, respectively, a species belongs. Colour codes are the same of (b), which show the boxplots of the *nmi* distribution between different categories.

## 4. Discussion

By applying network models to a long-term series of captures of spring migratory passerines at a stopover site, we showed that changes in the timing of migration of African-European landbirds lead to non-random patterns in the assemblage of co-occurring migrants. We found that modularity is a common pattern in temporal co-occurrence networks, with a trend for an increase over the years. This indicates a general tendency of birds to arrive at the stopover area in well-defined groups of species characterized by common migration phenology, a tendency likely to be influenced by changes in climatic conditions and to which species respond by restructuring their migration patterns in a non-random way.

Spring migration phenology of birds crossing the Mediterranean Sea can be related to specific ecological factors associated with travel distance and competition for nesting sites, as well as to the degree of sexual dimorphism [55]. For instance, species wintering closer to the breeding areas (e.g., North Africa) usually have faster adaptive responses due to their ability to better track changing conditions at the breeding grounds [56]. Latitudinal differences in wintering sites might thus determine the phenology of spring migration because species wintering north of Sahara advance their peak date of passage of more days compared to sub-Saharan ones [20]. Moreover, recent findings have shown a substantial increase (up to one day per year) of the migration window for species wintering south of the Sahara Desert, possibly related to more favourable conditions at wintering areas that allow faster refuelling, earlier departure, and/or longer residence time [20]. The strong modular subdivision observed over the study period (with the only exception of 2010) suggests that passerine species moving within the African-West Eurasian flyway during spring migration always followed a well-defined order of migration, a tendency likely to be predominant in those species forced to cover larger distance to reach their breeding areas.

How migratory birds manage to detect and adapt to changing environmental conditions during their journey to breeding areas remains an open question, possibly related to whether shifts in migration phenology are driven by microevolutionary changes or phenotypic plasticity [57]. Phenotypic plasticity may provide faster adaptive responses, especially for species travelling shorter distance and capable of immediate response to changing environmental conditions [58]. Although our findings do not allow for direct evidence, they nevertheless indicate a close relationship between bird’s phenotypic responses to migration and co-migration fidelity. This would explain why, for instance, North-African migrants showed lower co-migration fidelity than trans-Saharan ones, a pattern likely to be related to more flexible responses to changing climate conditions and possibly influencing the repeatability and synchrony of migration phenology over time.

### 4.1 Consequences for coexistence

Although co-occurrence patterns cannot be used to infer actual interspecific interactions (see [38] and references therein), they can reveal potential for interferences among species, especially when these are forced to share small areas in limited time frames, such as at migratory bottlenecks [59]. This is the case of stopover areas, where birds stop to rest, refuel and seek shelter during their migration activity, and where the decision on where, when and how long to rest is driven by physiological adaptations associated with migratory behaviour and habitat quality [60–62]. For instance, the amount of forest cover and productivity relates to the two-step mechanism determining the spatiotemporal use of stopover areas, *via* identification of high-quality habitat prior to landing and sustained refuelling rates during stopover [61,63]. Overall, stopover areas are important to replenish energetic reserves and can influence subsequent flight ranges [64,65]. However, stopover sites can have functions different from providing food, and their ecological context (e.g., proximity to ecological barriers, spatial isolation) and intrinsic characteristics (e.g., diversity and abundance of resources) may determine the use made by migrants [20,62,66,67].

Most individuals do not use Ponza for refuelling [67], as they might just need to rest before continuing the last part of their journey [68,69]. Therefore, coexistence at stopover areas might not be regulated exclusively by avoidance of competition for resources, but also by the need to reduce the risk of predation or to gain information on novel habitats and resources [33,34,70]. Almost all species investigated in our study have a strictly carnivorous diet and feed mainly on insects, with the exception of the European turtle dove, *Streptopelia turtur* and the Eurasian golden oriole *Oriolus oriolus*, which are classified mainly as granivores’ ground feeding species, but have nevertheless variable foraging niches [53]. Several studies have shown that species with similar diets switch to alternative foraging substrates and different predatory behaviour to avoid competition [53,71]. Our findings show no association between co-migration fidelity and foraging strategies, suggesting the existence of species-independent foraging mechanisms promoting coexistence at stopover areas.

However, if refuelling is not the main reason for migrants to land in an area [20], social (facilitative) interactions could explain the observed differences in species co-migration fidelity. Social interactions have been observed in migrating songbirds that use interspecific calls as cues to estimate stopover quality [33]. More recently, Gayk et al. (2021) [72] showed how new world warblers that migrate in mixed-species flocks produce flight calls to reduce disorientation during migration or to increase the chance of finding high quality stopover sites, a pattern even more pronounced in phylogenetically close species that breed at the same latitude with overlapping migration timing. Thus, it could be hypothesized that the observed tendency to migrate in well-defined and consistent subgroups of species with synchronized migration times is functional to optimize the costs associated with predation risks *en route* and resting at stopover area, in order to improve their competitive ability for territories at breeding areas.

## 5. Conclusions

As animal migrations involve the simultaneous movements of individuals and species in space and time, the emergence of potential interactions is far from being rare and is likely to influence the costs and benefits of migration. Long-term phenological changes have the potential to determine when, how long and with whom each species co-occurs with other members of the community. Furthermore, the consistency and synchronization of such changes might influence the global economy of migration. Here we show that modularity is a common pattern in the multi-species assemblage of African-European migratory landbirds, meaning that during spring migration birds follow a well-defined schedule driven by common responses to the effects of climate change on the timing of migration. Such pattern is likely to increase over time, mainly affecting species with low phenotypic plasticity and which cover longer distance during migration, with consequent fragmentation of migration timing. Further studies are however needed to better understand the consistency of observed patterns and the consequence for coexistence, for instance by extending the spatiotemporal coverage of analysis to species’ co-occurrence at wintering, stopover and breeding areas, or by experimental tests of the effects of competitive interactions in the field. Our study represents a first attempt in modelling multi-species temporal data to gain new insights in the consequence of long-term changes in migration phenology on the restructuring of avian assemblages *en route*.

## Acknowledgements

We would love to dedicate this work to the loving memory of our friend and colleague Prof. Dario Angeletti, torn from life too early. We would also like to thank Prof. Hanna Kokko for the stimulating discussion that greatly improved the drafting of the manuscript. The authors would also like to thank all the hundreds of volunteers at the bird ringing station of Ponza (http://www.inanellamentoponza.it), who have made it possible to constantly collect data over the years. This is publication N. XX of the *Piccole Isole* project.

## Conflict of Interest

The authors have no conflict of interest to declare.

## Authors’ contributions

BB conceived the paper together with CC; BB analysed the data and led the writing of the manuscript; MC prepared and analysed the data. IM and LF contributed to data collection and to writing of the manuscript. All authors have no perceived conflict of interest and contributed critically to the writing of the manuscript and gave final approval for publication.

## Funding statement

Data used for this study were collected thanks to funds of the University of Ferrara, University of Veterinary Medicine, Vienna, Grant Nr. 196451/V40 of the Research Council of Norway, Grant Nr. P31037-B29 of the Austrian Science Fund (FWF), and a number of donations and contributions to Centro Italiano per lo Studio e la Conservazione dell’Ambiente (CISCA).

## Data Availability Statement

Data used in this paper are available at Figshare:

Bird ringing data: https://doi.org/10.6084/m9.figshare.20438997.v1

Selected bird traits: https://doi.org/10.6084/m9.figshare.20439027.v1

R code: https://doi.org/10.6084/m9.figshare.20438982.v1.

Aggregated data-only can also be found at https://phaidra.vetmeduni.ac.at/view/o:114.

## Notes

https://phaidra.vetmeduni.ac.at/view/o:114

https://doi.org/10.6084/m9.figshare.20438997.v1

https://doi.org/10.6084/m9.figshare.20439027.v1

https://doi.org/10.6084/m9.figshare.20438982.v1

### Competing Interest Statement

The authors have declared no competing interest.

### Summary of Updates

After a first round of revisions, the original paper has been drastically changed following the suggestions provided by expert in the fields. We therefore changed the focus of the original submission by focusing on the whole avian assemblages, without distinguish between males and females and focusing on one main structural properties of the networks, modularity. A lot of changes have been made throughout the text.

