## Supporting Information for "Modularity in temporal co-occurrence networks at a stopover site reveals strong co-migration fidelity in African-European migratory landbirds over time" (from another version of this preprint: version 1, 2022-02-19)

**Sex differences in migration timing influence the network structure of avian assemblages at a stopover site**

^*^Corresponding author

**Figure S1** – (a) Temporal trend of the total number of caught individuals for the 17 species selected in this study. Red and blue lines are for females and males, respectively. (b) Boxplots showing the distribution of the number of individuals for each species, with red and blue boxes for females and males, respectively.

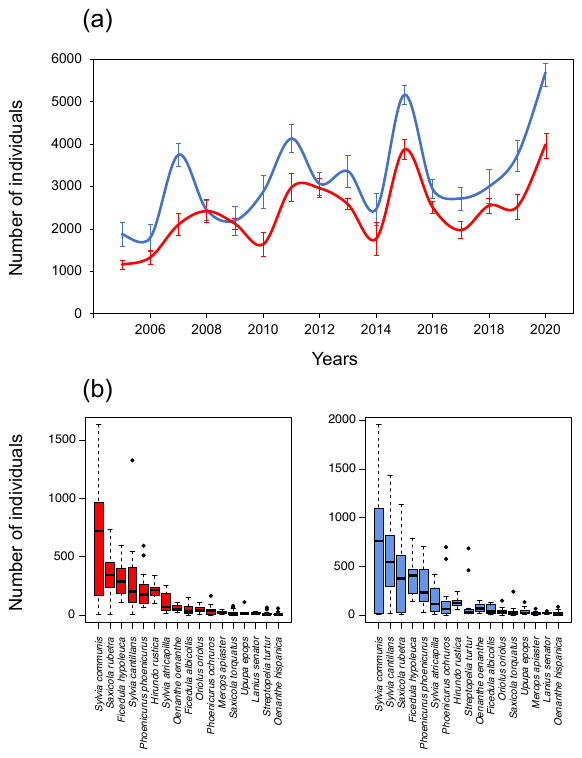
